# Fully Automated EEG Source Imaging Using Structured Sparsity for Single and Multiple Synchronous Epileptic Activities

**DOI:** 10.1101/2025.11.04.686464

**Authors:** M. Aud’hui, A. Kachenoura, L. Albera, I. Merlet, M. Yochum, P. Van Bogaert, M. Kuchenbuch, P. Benquet

## Abstract

Accurate localization of epileptic zones from High-Resolution ElectroEncephaloGraphy (HR-EEG) data can be challenging, especially when multiple synchronous zones are involved, and is highly dependent on the chosen EEG Source Imaging (ESI) method. Since a given scalp-level electrical pattern can result from multiple source configurations, ESI methods address this ill-posed inverse problem by imposing constraints on the structure of underlying sources.

Here, we present an efficient approach that imposes sparsity on both the source-level activity and its spatial gradient. Unlike other methods that generally require a heuristic choice of a regularization parameter that balances between data fidelity and constraints, our method iteratively adjusts the parameter value based on the noise level in a fully automated way.

The performance of the new method is evaluated across different scenarios of realistic synthetic HR-EEG data, including unifocal and synchronous multifocal cortical epileptic activity. Based on multiple performance indices, we demonstrate that the proposed approach outperforms traditional methods in accurately reconstructing epileptic sources. We also show that the method reduces polarity artifacts responsible for ghost sources and spatial discontinuities. Its ability to recover homogeneous, well-delineated regions of activity is further confirmed using real EEG data capturing a typical absence seizure.

## 1. Introduction

ElectroEncephaloGraphy (EEG) is a key tool in the identification of epilepsy, as it detects brain abnormal interictal and ictal signals, which are essential for the electro-clinical classification of epilepsy syndromes. It offers notable advantages over other neuroimaging modalities, including its non-invasiveness and remarkable temporal resolution. With the advancement of High-Resolution EEG (HR-EEG), which utilizes up to 256 electrodes, it has become possible to generate source-level activity maps with spatial resolutions comparable to MagnetoEncephaloGraphy (MEG) (Hedrich et al. 2017). Indeed, HR-EEG offers superior spatial resolution compared to standard EEG, enabling more precise localization of epileptic foci. For instance, a recent study reported successfully mapping the epileptogenic network in pediatric and young adult patients with drug-resistant epilepsy undergoing surgery, using non-invasive interictal HR-EEG data in different subtypes of focal epilepsies (Corona et al. 2023).

Unifocal cortical epileptic activities are commonly observed in various pathologies like focal cortical dysplasia, heterotopia, tumors, stroke, trauma and vascular malformations (Nascimento et al. 2023), and precise localization of the Epileptic Zone (EZ) is crucial for evaluating the feasibility of neurosurgical intervention. In this context, HR-EEG-based source localization could provide valuable information to guide invasive stereotaxic intracranial EEG recordings (sEEG), which can be essential in the next step for precisely delineating the EZ, depending on the clinical context (Li et al. 2022). Despite methodological improvements, including the use of the cortex as source space, the analyses in individual space, the exploitation of the wide frequency bands and of realistic head modeling, many challenges remain in achieving accurate source localizations. These challenges are even more pronounced when epileptic activity is multifocal or involves a large cortical network. This is the case in developmental and epileptic encephalopathy, tuberous sclerosis, extensive polymicrogyria, or generalized genetic epilepsies like absence epilepsies. In such cases, while surgical intervention is generally ruled out, identifying the underlying cortical network of epileptic activity can nevertheless be useful for scientific purposes, aiding in the understanding and classification of epilepsy syndromes. However, the accuracy of source localization in these contexts still needs to be thoroughly evaluated.

Source reconstruction and localization are highly dependent on the used EEG Source Imaging (ESI) method to solve the inherent EEG inverse problem. ESI methods can be classified into two categories: i) the equivalent source dipole methods, which consider that isolated dipoles represent the brain sources with unknown positions, and ii) the distributed sources techniques where each brain source is represented by a large number of dipoles distributed in the whole cortical surface. It is well accepted that the distributed methods are more physiologically adapted for characterizing epileptic regions. Among the ESI distributed methods, the well-known weighted Minimum Norm Estimates (wMNE) is the most widely used solution. This algorithm originates from the MNE (Hämäläinen and Ilmoniemi 1994) which exploits the L2-norm as a regularizer to find the solution that best describes the measurements in a least-squares sense. However, contrary to the classical MNE which tends to favor superficial sources, wMNE proposes to circumvent this bias by imposing a predefined weight on each source dependent on its distance from the scalp electrodes. Another variant of wMNE, which is widely adopted by the neuroscience community, is the popular standardized Low-Resolution Electromagnetic Tomography (sLORETA) (Pascual-Marqui 2002). The idea of the sLORETA algorithm consists in normalizing the MNE solution according to the variance of each estimated source. Even if these popular methods robustly target the EZ, they suffer from significant limitations. Indeed, as they promote a smooth spatial distribution, they generally lead to over-smoothed (blurred) solutions, and are not able to distinguish between the close patches. This can be very problematic if we want to estimate specific EEG metrics such as functional/effective connectivity at the cortical surface (source space). To cope with these aforementioned limitations, our group proposed in (Becker et al. 2017) a new method, called Source Imaging based on Structured Sparsity (SISSY). This algorithm includes a L1-norm regularization term, which imposes sparsity on the estimated source distribution. Provided that the hyperparameters are correctly tuned, we demonstrated that SISSY can separate even close sources and avoids amplitude-biased source estimates. Indeed, the choice of the regularization parameter value is critical in regularization-based methods and a major difficulty lies in its determination. Thus, in this study, we present a new version of SISSY named APESISSY (Adaptive Parameter Estimation SISSY), where the regularization parameter that balances between the amount of data fidelity and the regularization term is identified automatically.

To analyze the performance of the APESISSY algorithm, we conducted an extensive simulation study with highly realistic EEG data. Indeed, assessing ESI methods based on simulated data is convenient, as the scalp EEG is derived from a known source-level activity map which constitutes the ground truth. In this paper, five scenarios were designed, involving spike-wave activity, the EEG signature of absence epilepsy, in one (equivalent to focal epilepsy) to twelve cortical regions of various extent. Simulations were generated with a virtual human brain model (eCOALIA) (Wendling et al. 2024; Köksal-Ersöz et al. 2024), based on interconnected neural masses respecting the physiology and anatomy of the brain, able to provide realistic clinical scalp EEGs (Maliia et al. 2024). Contrary to the SISSY paper where the used number of scalp EEG electrodes was limited to 91, the quantitative evaluation of the proposed method is conducted on scalp epileptic HR-EEG signals (256 electrodes). A performance comparison study is also conducted with the two most widely used ESI methods, namely wMNE and sLORETA. Eventually, we also compared and analyzed the results obtained with the three ESI methods on real HR-EEG data recorded during a seizure from a patient with diagnosed Childhood Absence Epilepsy (CAE).

## 2. Methods

The pipeline (Fig. 1) involves the simulation of source activity (ground truth), the generation of the EEG at scalp level, the estimation of source activity with the three ESI methods and the assessment of their performance with statistical indices.

**Figure 1:**
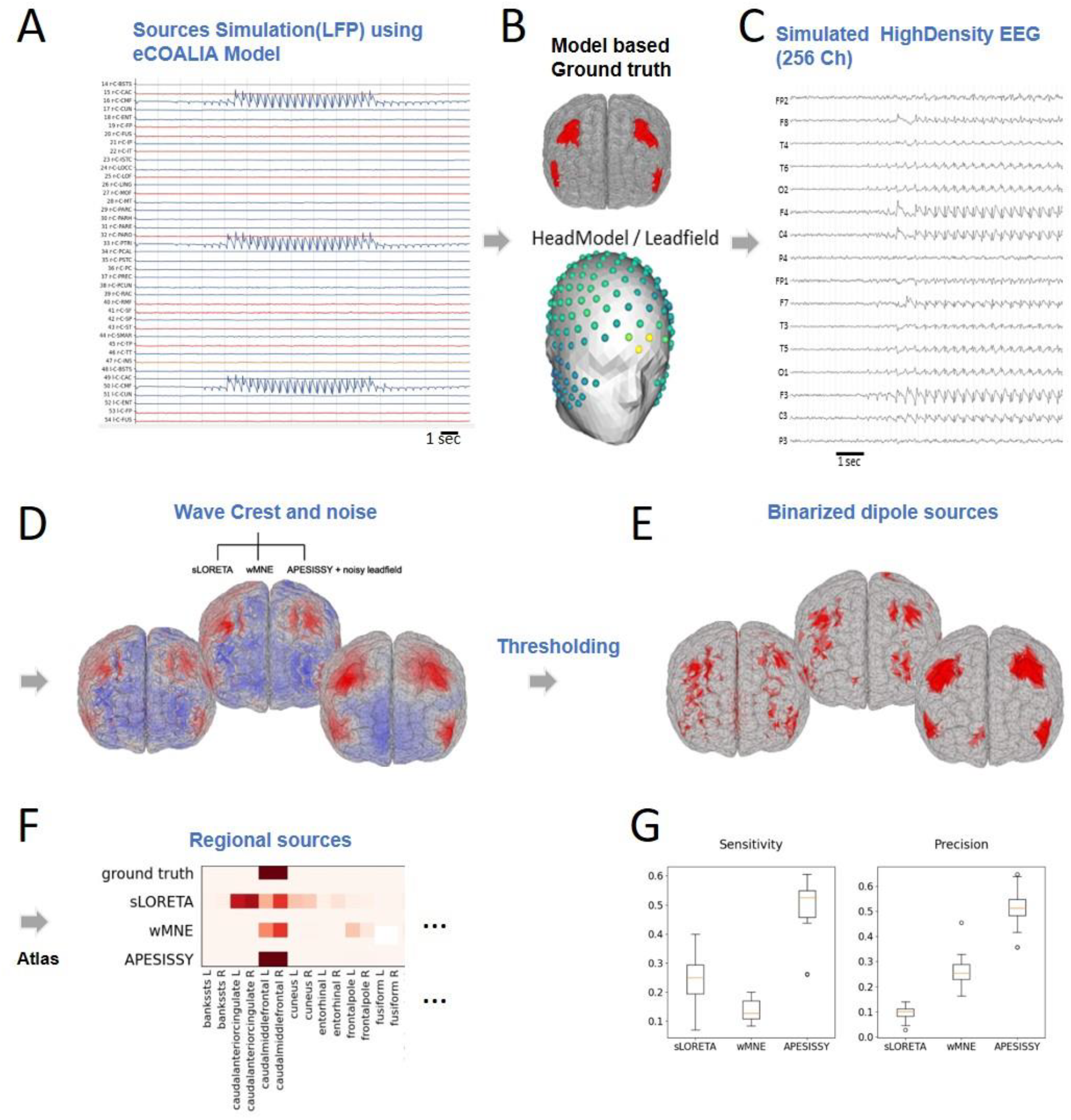
Pipeline and source localization methods comparison based on the simulated HD-EEG signals. **A.** Generation of multiple spike-wave discharges in one or multiple cortical areas as ground truth using eCOALIA. Source-level activity is simulated at the regional level with interconnected neural masses and mapped to individual dipoles using a template-based atlas. **B.** This scenario of model based ground truth is projected onto virtual surface electrodes using a lead field matrix. **C.** 256-channel EEG is then generated from which we extract a noise period. **D.** Instantaneous signal of interest at the crest of a wave are **E.** subsequently binarized through a thresholding step. **F.** Regional sources localized with the various source localization methods are finally compared with the known simulated ground truth to **G.** derive various metrics for assessment.

### 2.1. Forward modeling

We used Brainstorm (Tadel et al. 2011) to derive a 15,002-vertex cortex mesh from a reference anatomy (ICBM 152 2023). Dipoles were mapped to 68 Regions Of Interest (ROIs) according to the Desikan-Killiany atlas (Desikan et al. 2006). An oriented lead field matrix of size 15,002×256 was computed with OpenMeeg implemented in Brainstorm (Gramfort et al. 2010) using the Boundary Element Method (BEM) with a 3-layer model comprising the brain, skull and scalp (conductivities 1, 0.0125 and 1 reciprocally). This lead field matrix was used to solve the direct problem. To avoid committing the « inverse crime » (Wirgin 2004), a second lead field matrix was derived from the first and dedicated to solving the inverse problem. It was obtained by multiplying its terms by those of a randomly generated Gaussian distribution of mean 1 and standard deviation of 0.05.

### 1.1. Simulations

The temporal dynamics of the epileptic activity of each dipole were simulated using the whole brain model named eCOALIA based on human brain anatomy and connectivity, described in detail in (Wendling et al. 2024; Maliia et al. 2024; Köksal-Ersöz et al. 2024). The layered Neural Mass Model (NMM) is designed to reproduce the dynamics of the average activity of a neuronal population consisting of synaptically connected neuronal subpopulations of glutamatergic excitatory neurons and inhibitory GABAergic Interneurons (INs). The model considers two types of GABAergic INs (parvalbumin positive cells, PV+ INs; somatostatin positive cells, SST+ INs), and takes into account the positions of synapses across layers to compute the resulting Local Field Potential (LFP) (Köksal-Ersöz et al. 2022; Wendling et al. 2024). In brief, according to the NMM formalism, in each subpopulation, a dynamic linear transfer function converts presynaptic information (i.e. the average pulse density of afferent action potentials) into an average membrane PostSynaptic Potential (PSP), which can be either Excitatory (EPSP) or Inhibitory (IPSP). The kinetics (rise and decay times) of PSPs reproduce those of real PSPs. In turn, a static nonlinear function (sigmoid shape) relates the average membrane potential of a given subpopulation to an average pulse density representing the sum of action potentials fired by neurons. Neural masses are connected through long-range glutamatergic projections among pyramidal neurons and GABAergic interneurons (Wendling et al. 2024). At the global level, the large-scale model is constructed based on 66 regions of interest (ROIs) from the standard anatomical parcellation of the Desikan-Killiany atlas. The right and left insula were excluded, leaving 66 brain regions. Each neural mass output in this case represents the LFP of one atlas region, where the activity is assumed to be homogeneous.

Three scenarios were simulated, involving from one to twelve synchronous epileptic regions. In all scenarios, cortical spike wave discharge was triggered by a feedforward excitation resulting from a thalamic input. The thalamic activity was induced by an electrical stimulation. Depending on the scenario, each neural mass had its own sets of parameters regulating the activity of the pyramidal cells and the GABAergic neuronal populations. For each scenario, we slightly modified the gain parameter of the pyramidal cells of each region by adding a value taken from a Gaussian distribution of mean zero and standard deviation 0.5 to obtain a panel of 20 Monte Carlo (MC) instances of source-level activity. At the source level, the activity was then mapped to all 15002 dipoles using the mesh-derived atlas, and a 256-channel EEG was generated through multiplication with the lead field matrix.

Epileptic regions in the three scenarios were respectively:

1. one cortical area: right Caudal Middle Frontal gyrus (CMF)
2. four cortical areas: CMF and Pars TRIangularis (PTRI) (both bilateral)
3. 12 cortical areas: CMF, Inferior Parietal cortex (IP), PreCUNeus (PCUN), PoSTerior Cingulate cortex (PSTC), Superior Frontal cortex (SF) and SupraMARginal gyrus (SMAR) (all bilateral)

Scenarios 1 and 2 were designed to simulate focal activity and moderate multifocal activity respectively. The third scenario was explicitly designed to mimic a real absence seizure, and regions were selected to match as closely as possible the regions associated the most often with absence seizures in functional Magnetic Resonance Imaging (fMRI) (Moeller et al. 2008; Li et al. 2009; Bai et al. 2010; Berman et al. 2010; Masterton et al. 2013; Carney and Jackson 2014) and MEG studies (Westmijse et al. 2009; Tenney et al. 2013; Miao et al. 2014; Wu et al. 2017; Gadad et al. 2018; Wang et al. 2023). This modeling comes with its limitations due in part to the anatomical constraints by the atlas. In particular, IP and SMAR were selected to account for the well-documented involvement of the temporo-parieto-occipital junction in absence seizures (Tang et al. 2016; Wu et al. 2017). The resulting EEG can be seen with a transversal montage in Fig. 2. Two additional sub-scenarios were derived from the third one (scenario 3bis and scenario 3ter). In scenario 3bis, all epileptic regions were reduced to smaller subregions of 100 dipoles, while in scenario 3ter, they were reduced further to subregions of 20 dipoles. These sub-scenarios were designed to study the sensitivity of the methods to changes in the extent of the involved regions.

**Figure 2.**
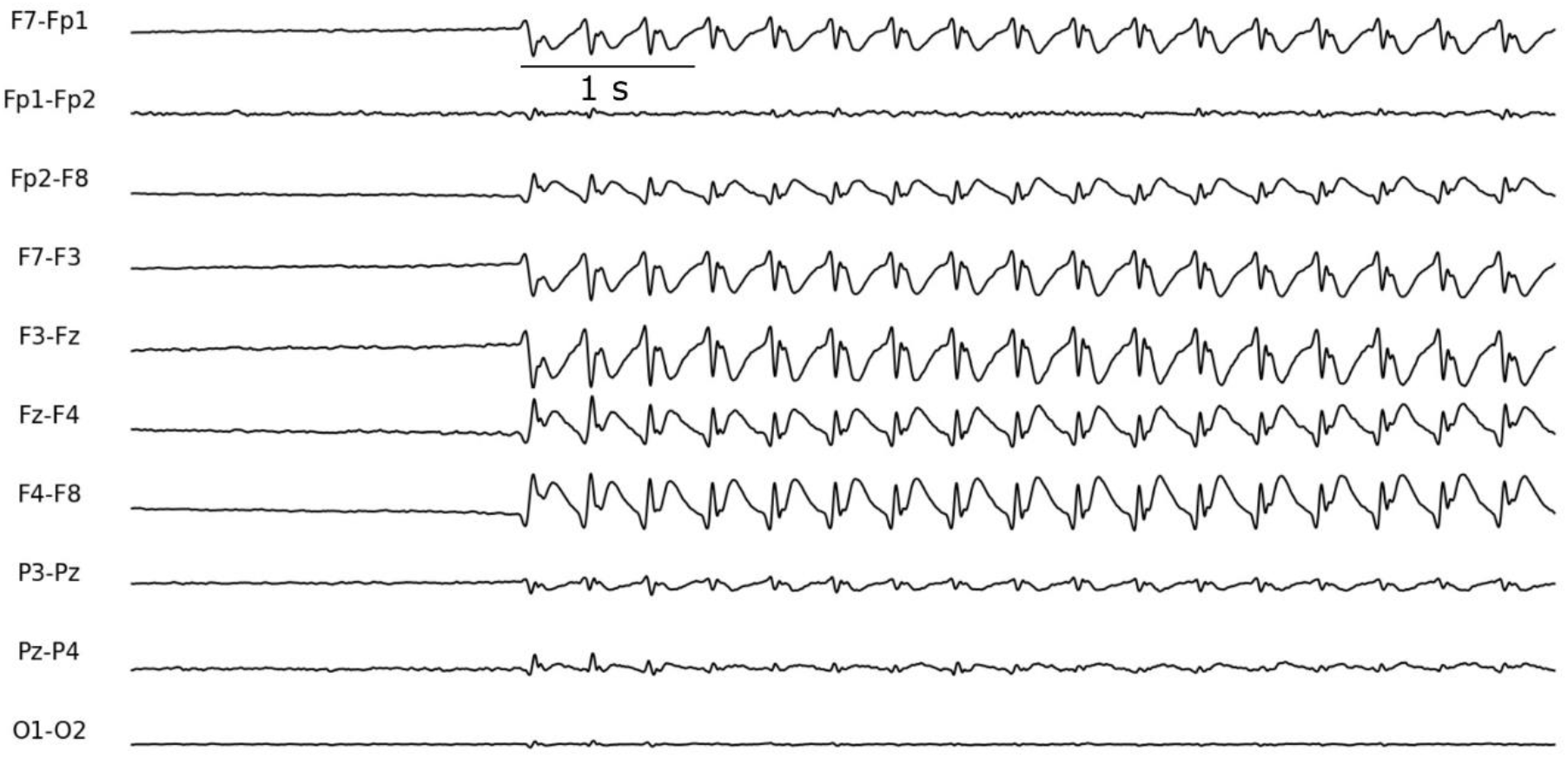
Scalp EEG signal generated from a simulated absence seizure involving the regions defined in scenario 3, displayed with transversal montage.

### 1.2. Source computation

As stated before, we have shown in our previous work (Becker et al. 2017) that the SISSY algorithm exhibits better performances in separating multiple epileptic sources than classical ESI methods. In this section, we present an updated version of SISSY, named APESISSY, where we propose an automatic identification of one of its penalty parameters. A natural approach to solve the ill-posed EEG inverse problem consists in finding the solution that best describes the measurements in a least squares sense. In the presence of noise, this is generally achieved by minimizing the following cost function:

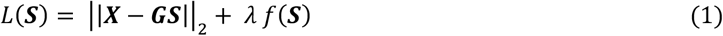

where ***X*** is the scalp EEG signal, ***G*** is the lead field matrix computed from a patient-specific brain mesh comprising a number of dipoles dispatched along the cortex, ***S*** is the sources solution, i.e. the current density values of the dipoles. The first term on the right-hand side of (1) is generally referred to as the data fit term and characterizes the difference between the measurements and the surface data reconstructed from given sources. The second term is a regularization term and incorporates additional constraints on the sources according to a specific prior information. The regularization parameter *λ* is used to manage a trade-off between data fit and a priori knowledge, and depends on the noise level, since the gap between measured and reconstructed data is expected to become larger as the Signal-to-Noise Ratio (SNR) decreases. For all scenarios considered in this paper, we performed source computation on the instantaneous signal extracted from the crest of a wave, when the signal is strong and steadier than at the top of the spike. Consequently, the cost function (1) to be minimized can be reformulated as follows:

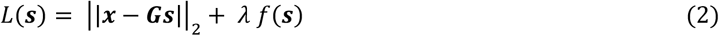

where ***x*** and ***s*** are the scalp EEG and brain source vectors at the time corresponding to the crest of the wave.

Before presenting in details the APESISSY, let’s briefly recall the principal of SISSY algorithm. This latter is based on the hypothesis of the sparsity of both the sources and their spatial gradient. Thus, its regularization term relies on the L1 norm or summation of absolute values, applied both to ***s*** and to ***Ts***, where ***T*** is a variational operator computing the differences between pairs of adjacent dipoles. The regularization term *f*(***s***) can then be expressed as ||***Ts***||_1_+ α||***s***||_1_, and leads to the following minimization problem:

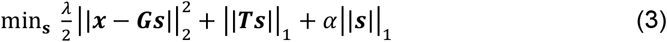

where the penalty parameter α permits the adjustment of the size of the reconstructed source region (the larger α, the smaller the estimated source). The regularization parameter *λ* balances between the reconstruction error and the spatial distribution constraint. Compared to the L2 norm, the L1 norm exerts a lesser constraint on extreme values and favors the sparsity of the solution, i.e. the maximization of the number of null instances. Due to the minimization of the L1 norm of the spatial gradient, SISSY is expected to enable to rebuild well delineated homogeneous patches of activity. Note that in the SISSY approach, while parameter *α* is generally chosen small it is more challenging to fix the regularization parameter *λ*, since it depends on both the noise level and the number of EEG channels. Hence the interest in the APESISSY solution that we propose hereafter.

The APESISSY approach consists, therefore, in solving the minimization problem of (3) by computing a penalty parameter *λ*, which guarantees the solution that satisfy Morozov’s discrepancy principle (He et al. 2014). This is equivalent to solve the problem given by:

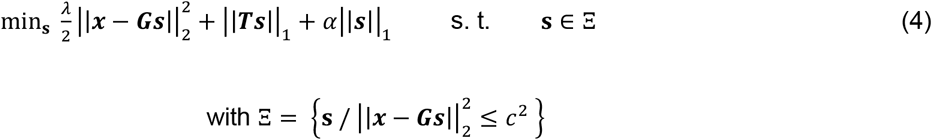

where *c* is a predetermined parameter, which is noise-dependent. Problem (4) can be solved by taking inspiration from the solution proposed to solve the TV-regularized image restoration problem about ten years ago (He et al. 2014). The solution is mainly based on the use of the Alternating Direction Method of Multipliers (ADMM) algorithm (Boyd et al. 2011). The latter is a splitting proximal optimization method, which overcomes the non-differentiability of the L1 norm and allows to achieve a closed form to update the penalty parameter in each iteration.

Let’s describe in more details the APESISSY approach based on the ADMM optimization strategy. First, let’s introduce additional variables related to the previous one through equality constraints:

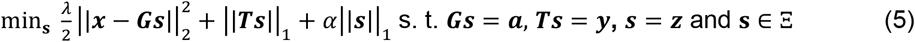

Then, problem (5) can be reformulated as the minimization of the following augmented Lagrangian function:

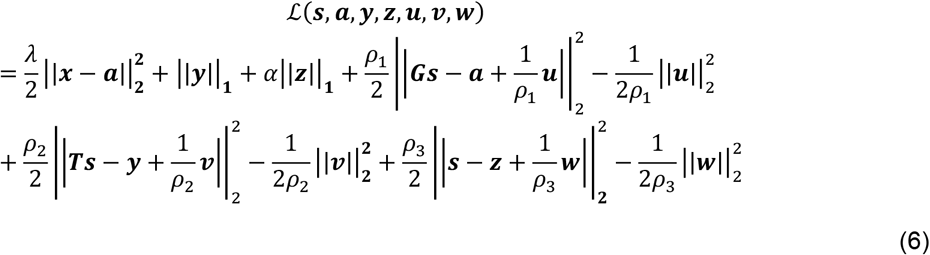

under the constraint **s** ∈ Ξ, where ***u, v*** and ***w*** denote the Lagrange multipliers. The main principle of ADMM is to alternate the minimization of the augmented Lagrangian function with respect to each variable by fixing the others. This leads to the following update rule of ***s***:

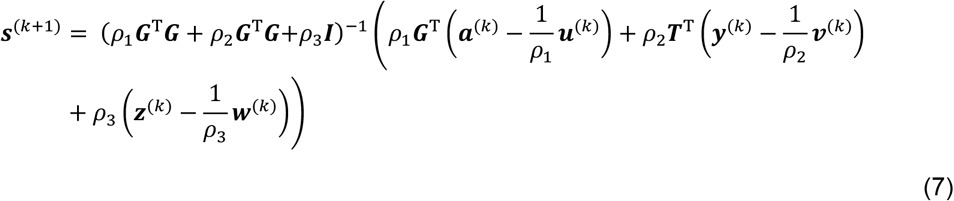

The minimization of the augmented Lagrangian function with respect to variable ***a*** by fixing the other variables gives the following update rule:

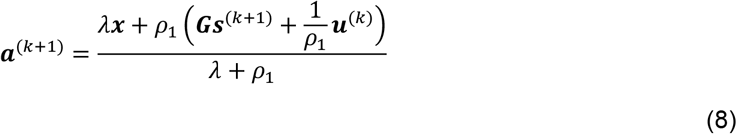

Then, the update *λ* ^(*k*+1)^ of the penalty parameter *λ* is derived by imposing Morozov’s discrepancy principle (Morozov, 1984). It requires to solve the following equality 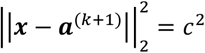 with respect to *λ* by injecting (8) in the previous equality, leading to the following solution:

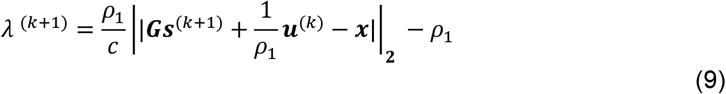

The penalty parameter *λ* is then replaced by *λ* ^(*k*+1)^ (9) in the definition of ***a***^(*k*+1)^ (8). The minimization of the augmented Lagrangian function with respect to variable ***y*** by fixing the other variables provides the following update rule:

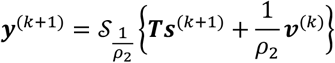

where 𝒮 is the soft-thresholding operator, defined by:

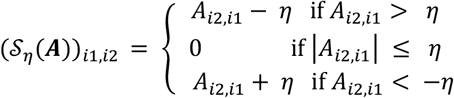

In the same way, we compute the update ***z***^(*k*+1)^:

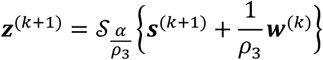

Finally, the Lagrange multipliers are updated by maximizing the Lagrangian function using a gradient ascent such that:

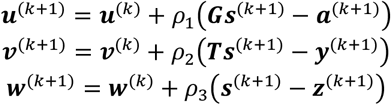

The parameters *ρ*_1_, *ρ*_2_ and *ρ*_3_ can be automatically tuned using an adaptive algorithm as proposed in (Boyd et al. 2011). The iterative APESISSY method is stopped using normalized difference between two consecutive estimations of ***s***:

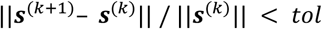

with *tol* fixed to 1e-3.

#### 1.3. Evaluation criteria

To assess the accuracy of the sources rebuilt by APESISSY and the other methods over the various scenarios, we used a set of four statistical metrics. From the computed dipole amplitudes, we first derived a binary array of activity; for each dipole, if its absolute amplitude exceeded 40 % of the maximum absolute amplitude among all dipoles, we set its activity to 1, otherwise we set it to 0. We defined a ground truth array of activity as well, composed of ‘1’s for dipoles pertaining to the “active” regions and ‘0’s for all other dipoles. From these two arrays, we measured four different metrics. The Sensitivity is defined as the number of dipoles active in both arrays (true positives) divided by the number of active dipoles in the ground truth (true positives + false negatives). The Precision is given by the number of dipoles active in both arrays (true positives) divided by the number of active dipoles in the first array (true positives + false positives).

We also computed the Dipole Localization Error (DLE) (Yao and Dewald 2005), to measure the similarity between the original and estimated sources configuration, defined by the formula:

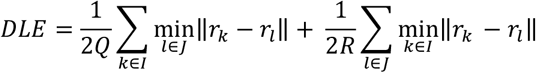

where *I* and *J* denote respectively the ground truth set and the estimated set of indices of the active dipoles, *Q* and *R* are the number of active dipoles in the ground truth and the estimated sources, and *r*_*k*_ the position of the k-th dipole.

Additionally, we chose to include a metric at the scale of the regions of the atlas, which is the most interesting scale for a large number of clinicians or researchers practicing source localization. To this end, for both ground truth and rebuilt sources, we computed a value of activity between 0 and 1 for each region by summing the activity of its dipoles and dividing the sum by the number of active dipoles in the region in the ground truth. We then divided these activities by the sum of activities over all regions to obtain distributions whose sums are equal to 1, and computed the Distance between these two Distributions (DD) as the average absolute difference between the distributed activities over the regions.

To assess the statistical significance of the obtained results, parametric (paired samples t-test) or non-parametric (Wilcoxon signed rank) tests were employed, contingent on the normality of the distributions of index values for all MC realizations, as determined by Shapiro tests. Statistical significance was determined with p < 0.05.

## 2. Results

The performance of the APESISSY algorithm is assessed on realistic simulated epileptic EEG data and compared to the two widely used approaches in the context of brain source localization, namely wMNE and sLORETA. These two methods were applied as recommended by one of the well-known open-source Matlab toolbox: Brainstorm (Tadel et al., 2011). As detailed in Section 2.1, we consider five scenarios with different numbers of epileptic patches and different patch sizes. In addition, all metrics were calculated on the 20 MC realizations, whatever the studied scenario. Finally, the behaviors of APESISSY were analyzed and compared to those of wMNE and sLORETA on real HR-EEG data from a patient diagnosed with typical CAE, to show the feasibility of such strategy in real life context.

### 2.1. Simulated data

#### 2.1.1. Scenario 1: single patch scenario

In this scenario, we simulated epileptic activity on a single region, namely the right CMF. The idea here is to assess the behavior of the methods in a simple case of distributed epileptic source. Figs. 3 A illustrates an example of the source imaging results obtained by the tested algorithms. We can see that the three methods exhibit a large activated area of positive polarity (red color) whose amplitude culminates in the simulated “active” region (right CMF). It is also interesting to see that both wMNE and sLORETA result on several patches of negative polarity (blue color) peripheric to the positive patch, while APESISSY gives only one such patch, more circumscribed and of lesser amplitude. Now, if we focus on brain maps of absolute activity after thresholding (Fig.3 A, bottom), APESISSY algorithm, which employs two sparsity constraints on the target sources, presents an excellent behavior in the case of a single source. Regarding wMNE and sLORETA, even though both methods target the desired region (right CMF), they lead to blurred source localization results and additional sub-regions far from the target.

**Figure 3.**
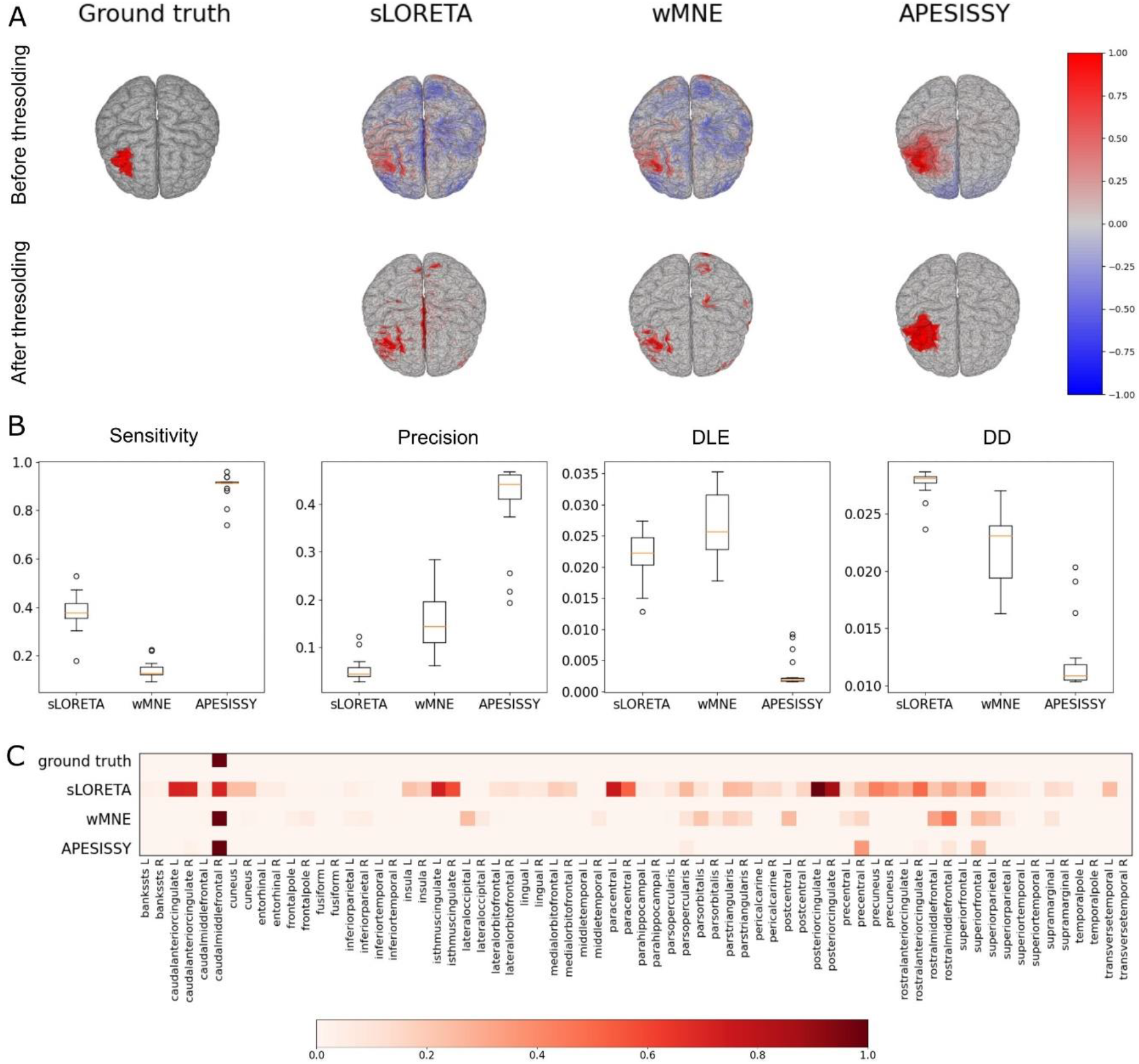
Source localization of epileptic-like activity simulated in a single cortical area (Scenario1). **A. Top.** Brain maps of simulated activity (ground truth) and of raw activity computed with the three methods, normalized by their maximum. Positively polarized activity appears in shades of red, and negatively polarized activity in shades of blue. **Bottom.** Brain maps of activity after thresholding. **B.** Boxplots displaying sensitivity, precision, DLE and distance between distributions of activity by regions, with all methods and for all 20 instances. **C.** Heatmap showing average normalized activity by region for ground truth and the three methods.

All the previous results are confirmed in Fig. 3 B, where we depict, for all compared methods the boxplots (over the 20 MC realizations) of the four metrics described in Section 2.3, namely Sensitivity, Precision, DLE and DD. It can be observed that APESISSY outperforms wMNE and sLORETA, for all considered metrics. Indeed, APESISSY performs much better than LORETA and wMNE in terms of Sensitivity and Precision (the medians are around 0.9, 0.4 and 0.1, respectively, for the Sensitivity, and 0.45, 0.05 and 0.15 for the Precision). This performance gap is blatant in the compared DLEs as well (under 0.005 for APESISSY, while sLORETA and wMNE are around 0.025), and in DDs of activity between regions (median is slightly over 0.01 for APESISSY, slightly over 0.025 for sLORETA and under 0.025 for wMNE). Another interesting visual result is given in Fig. 3 C, which displays the normalized activities of each region of the Desikan-Killiany atlas, averaged over the 20 MC realizations. As can be expected, the three methods capture the “active” region (CMF) but artefactually localize some additional regions as well. However, APESISSY, clearly, better fulfills the Ground truth. Indeed, sLORETA finds activity in particular in “deep” regions like cingulate cortices and in adjacent regions like the paracentral cortex, the SF or the rostral middle frontal gyrus. For wMNE, artefactual activity is more limited to adjacent regions, such as the rostral middle frontal gyrus, while it is further circumscribed for APESISSY to the right precentral and SF gyrus.

#### 2.1.2. Scenario 2

To deal with a multi-focal epilepsy context, the proposed model is evaluated on Scenario 2, which simulates the activity of four regions: bilateral CMF and bilateral PTRI. An example of source imaging result, before and after thresholding, is given in Fig. 4 A. All methods globally target the location of the four sources. In addition, as in scenario 1, wMNE and sLORETA exhibit a large artefactual negatively polarized patches, while APESISSY seems to be more robust to this polarity inversion. Indeed, APESISSY presents negative patches of limited extent and lesser amplitude. After thresholding, APESISSY maintains high performance and the estimated regions appear homogeneous and well-located. As in the single patch case, the remaining methods wMNE and sLORETA cannot accurately delimitate the spatial location and tend to give blurred and fragmented source localization results.

**Figure 4.**
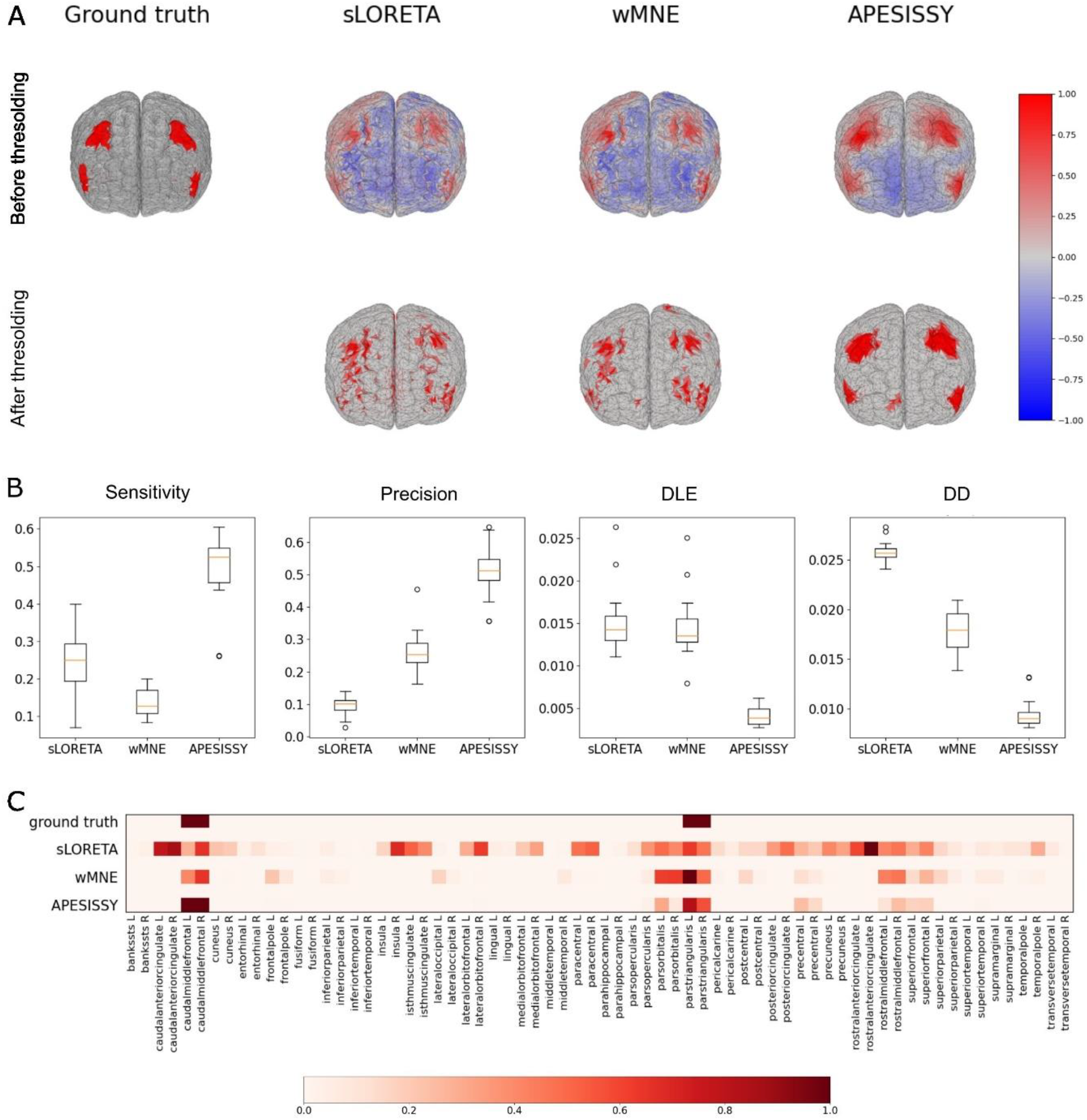
Source localization of epileptic-like activity simulated in 4 synchronous cortical areas (Scenario 2) **A. Top.** Brain maps of simulated activity (ground truth) and of raw activity computed with the three methods, normalized by their maximum. Positively polarized activity appears in shades of red, and negatively polarized activity in shades of blue. **Bottom.** Brain maps of activity after thresholding. **B.** Boxplots displaying sensitivity, precision, DLE and distance between distributions of activity by regions, with all methods and for all 20 instances. **C.** Heatmap showing average normalized activity by region for ground truth and the three methods.

Quantitatively, Fig. 4 B shows that all four metrics exhibit a performance gap between APESISSY and the two other methods. Median Sensitivity and Precision are both around 0.5 for APESISSY, while they are respectively under 0.3 and around 0.1 for sLORETA, and over 0.1 and around 0.25 for wMNE. The median DLE values for sLORETA and wMNE are three times higher than that of APESISSY, while the median DD is two to three times lower with APESISSY. Fig. 4 C illustrates the different regions of the Desikan-Killiany atlas that are located with the three methods. It confirms that all methods capture the four “active” regions. However, APESISSY shows only minimal artefactual activity in a limited number of adjacent regions. while the other methods find large extent activity in a number of artefactual adjacent (sLORETA and wMNE) or “deep” (sLORETA) regions.

#### 2.1.3. Scenario 3

In scenario 3, we simulated the activity of six bilateral regions: CMF, IP, PCUN, PSTC, SF and SMAR, for a total of 12 regions, six in the right hemisphere and six in the left one. Six patches (three in each hemisphere) appear on the ground truth map (Fig. 5 A). They correspond to the large CMF + SF frontal patches, the IP + SMAR patches, and the top of the PCUN patches. PSTC is a deep region and cannot be visualized with the view of Fig. 5 A. Globally, we observe that the reconstructed maps before thresholding correspond to the ground truth, particularly for APESISSY. After thresholding, activity in superficial active regions appears very sparse for sLORETA, and also for wMNE. APESISSY satisfactorily localizes the large frontal patch, as well as part of the right parietal patch, but introduces artefactual activity in the right superior parietal cortex. These results are reflected in the calculated metrics where a small Sensitivity is obtained for the three methods (Fig. 5 B), with a median peak over 0.25 for APESISSY, while sLORETA is under 0.15 and wMNE slightly over 0.05. Regarding the Precision, the obtained results are quite satisfactory, especially for APESISSY: around 0.6, 0.45 and 0.8 for wMNE, sLORETA and APESISSY, respectively. This means that, even if APESISSY does not target all the 12 regions, it does not mislocalize too many additional regions. Finally, contrary to the 2 first scenarios, the gap between APESISSY and the other methods narrows for the DLE, with all three medians falling within the 0.008 - 0.012 range, as well as for the DD metric (0.02, 0.018 and 0.015 respectively for sLORETA, wMNE and APESISSY). Fig.5 C gives more precise results on the regions that were well targeted or not. We see that the three methods succeed in the localization of the frontal bilateral CMF and SF but activity in bilateral PCUN, left IP and SMAR is overlooked. Activity in right IP and SMAR is overlooked as well with both sLORETA and wMNE. PSTC is detected by sLORETA only, though incorrectly completed by other regions of the cingulate cortex which are artefactual. Both wMNE and APESISSY detect activity in small left frontal pole, adjacent to SF. APESISSY detects artefactual activity both in the precentral and the paracentral gyrus as well, adjacent to CMF and SF, while sLORETA and wMNE detect activity in the rostral middle frontal gyrus, adjacent to both SF and CMF.

**Figure 5.**
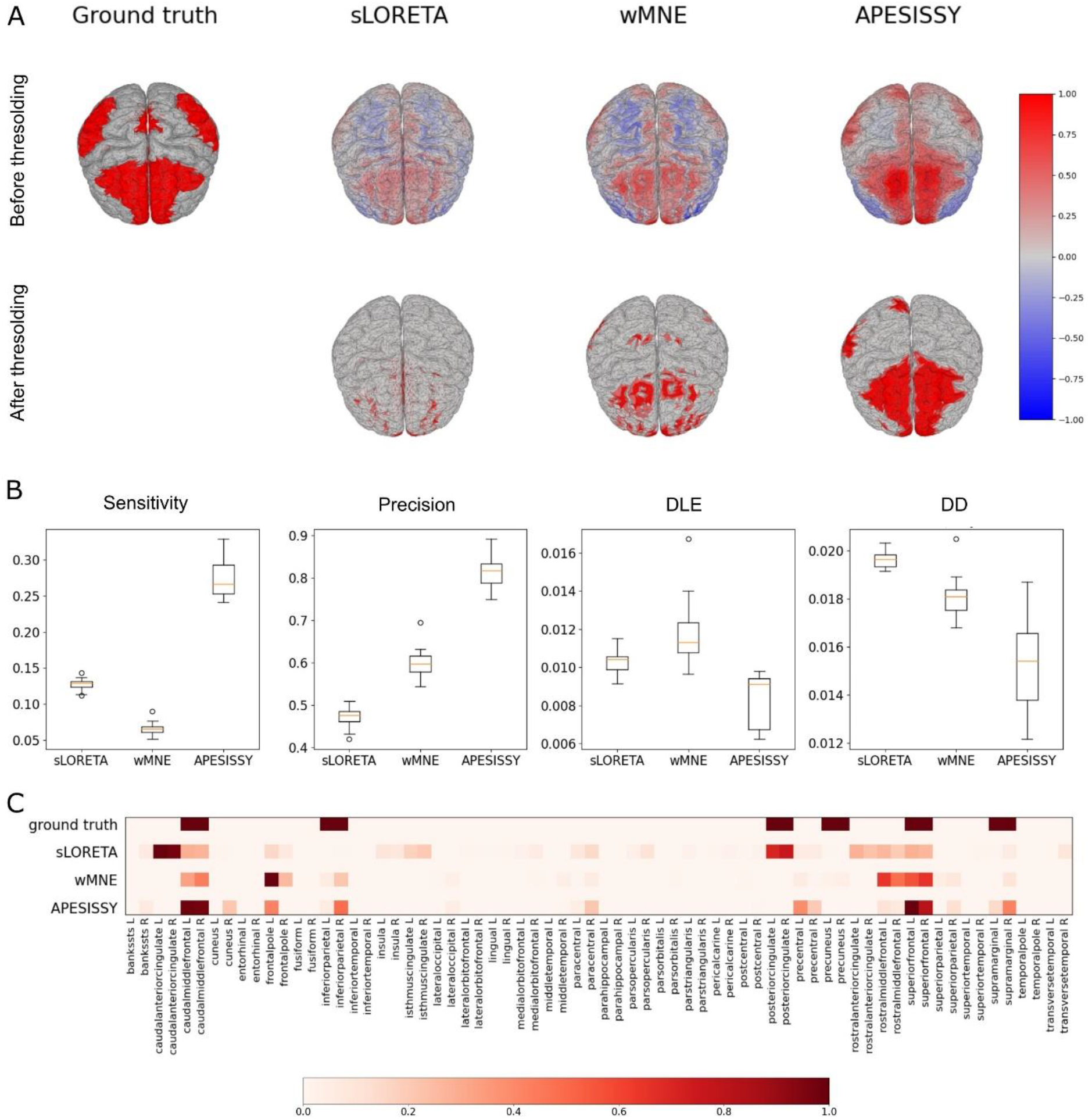
Source localization of epileptic-like activity simulated in 14 synchronous cortical areas (Scenario 3). **A. Top.** Brain maps of simulated activity (ground truth) and of raw activity computed with the three methods, normalized by their maximum. Positively polarized activity appears in shades of red, and negatively polarized activity in shades of blue. **Bottom.** Brain maps of activity after thresholding. **B.** Boxplots displaying sensitivity, precision, DLE and distance between distributions of activity by regions, with all methods and for all 20 instances. **C.** Heatmap showing average normalized activity by region for ground truth and the three methods.

#### 2.1.4. Scenario 3bis

Scenario 3bis involves the same “active” regions as scenario 3, but spike-wave activity is limited to a subset of 100 dipoles in each of these regions. We see (Fig. 6 A) that all superficial patches in the ground truth map are satisfactorily rebuilt with all methods. However, the CMF patches appear to have low relative amplitude, and APESISSY is the only method to restitute bilateral CMF activity after thresholding. Patches of negatively polarized activity are more circumscribed than in scenario 3. Median sensitivity (Fig. 6 B) is a little higher than in scenario 3, ranging from 0.15 for wMNE to around 0.3 for APESISSY. Precision increases for wMNE and APESISSY to values between 0.8 and 0.9, but only slightly to 0.2-0.25 for sLORETA. Median DLE is comparable for sLORETA and wMNE, around 0.009, and slightly lower for APESISSY, between 0.0075 and 0.008. DD criterion shows a slight gradual improvement from sLORETA to wMNE and APESISSY.

**Figure 6.**
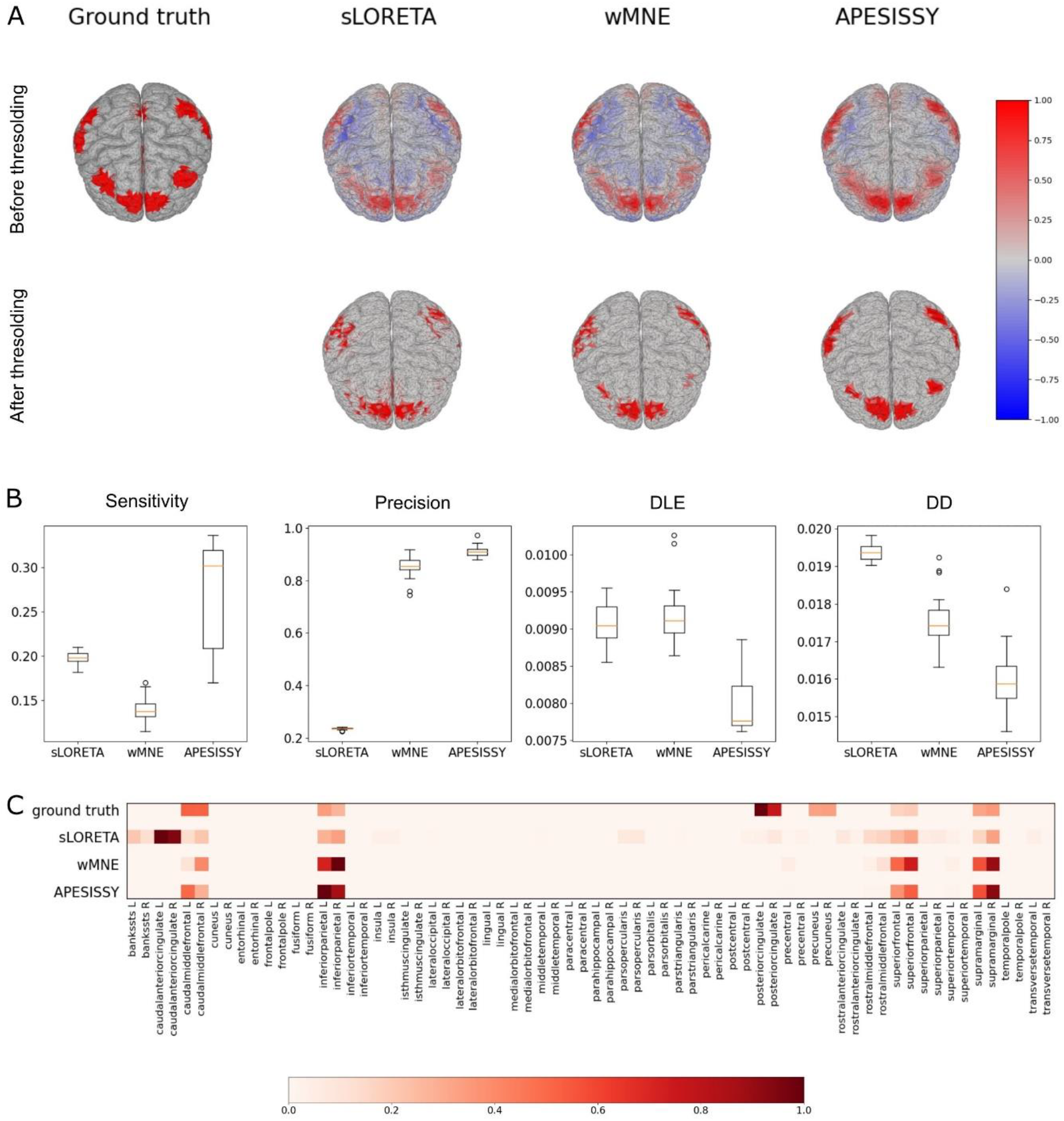
Source localization of epileptic-like activity simulated in 14 synchronous cortical areas of small extent (Scenario 3 bis). **A. Top.** Brain maps of simulated activity (ground truth) and of raw activity computed with the three methods, normalized by their maximum. Positively polarized activity appears in shades of red, and negatively polarized activity in shades of blue. **Bottom.** Brain maps of activity after thresholding. **B.** Boxplots displaying sensitivity, precision, DLE and distance between distributions of activity by regions, with all methods and for all 20 instances. **C.** Heatmap showing average normalized activity by region for ground truth and the three methods.

Of the 12 “active” regions, only the bilateral IP, SF, SMAR and the right CMF are well detected by all three methods (Fig. 6 C). Bilateral PCUN and PSTC are overlooked by all methods, and activity in the left CMF is only detected by APESISSY. sLORETA is the only method to detect significant artefactual activity in other regions, especially in the caudal anterior cingulate cortex and in the rostral middle frontal gyrus.

#### 2.1.5. Scenario 3ter

In scenario 3ter, the number of dipoles involved in spike-wave activity is further reduced to 20 in each “active” region. After thresholding, both wMNE and APESISSY provide a relatively accurate reconstruction of superficial epileptic region, but IP and SMAR patches are more clearly separated with APESISSY (Fig. 7 A). Patches of negative polarity gradually disappear from sLORETA to wMNE and APESISSY. As for all scenarios presented above, the solutions of wMNE and sLORETA are blurred, contrary to APESISSY which gives more circumscribed solutions. Compared to scenario 3bis, the Sensitivity (Fig. 7 B) continues to increase, especially for sLORETA and wMNE, ranging from 0.35 for wMNE to 0.45 for APESISSY. The precision of APESISSY and wMNE decreases to values around 0.6, and that of sLORETA to around 0.1. DLEs remain close, ranging from 0.0075-0.008 with APESISSY to slightly 0.0105 with sLORETA. The DD results are also similar to scenario 3bis, with a decrease for wMNE, resulting in slightly inferior values than APESISSY, though not significantly different (p-value of 0.62). Except for PCUN and PSTC, all regions are well localized by the three methods (Fig. 7 C). We have the same artefactual activity in other regions for sLORETA as in scenario 3bis.

**Figure 7.**
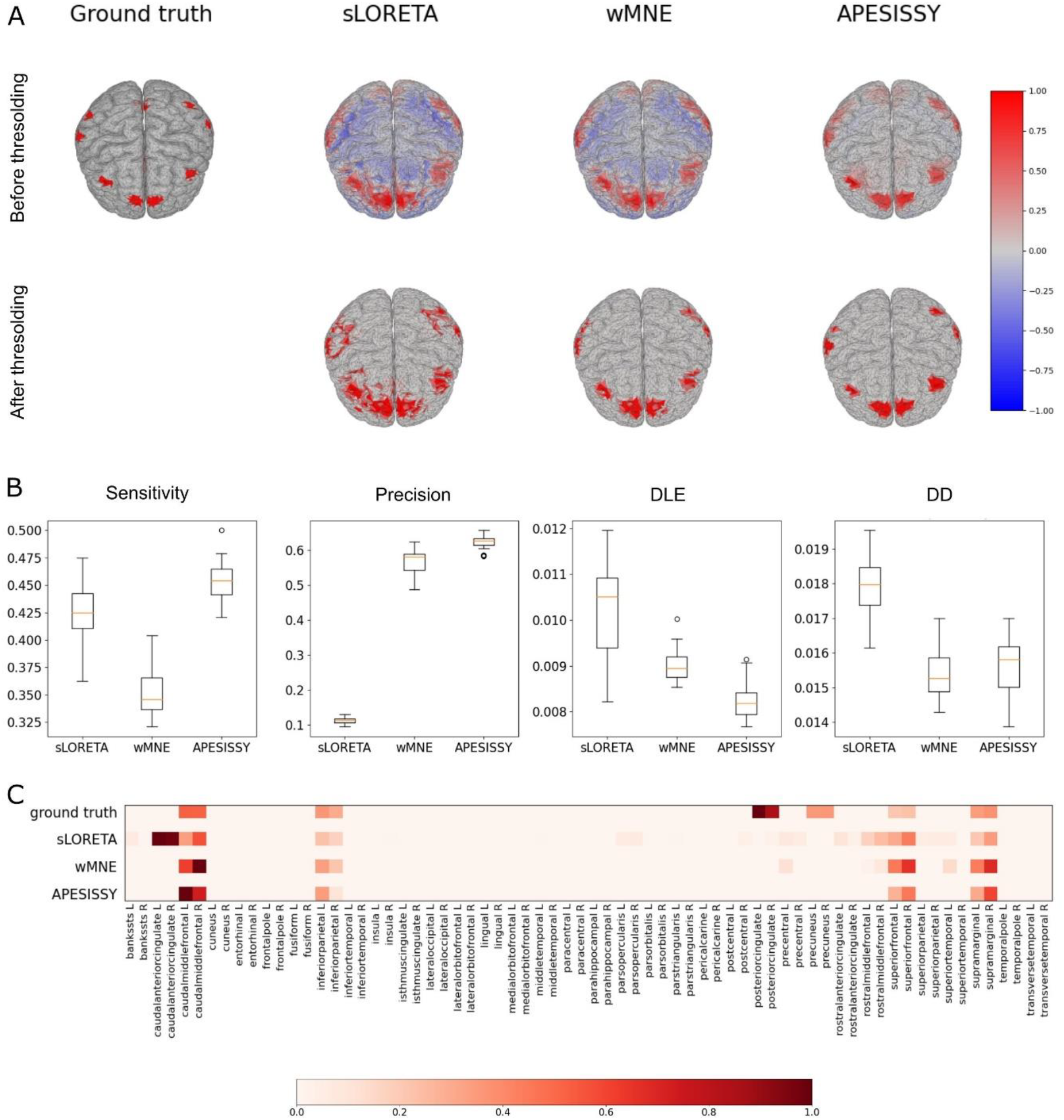
Source localization of epileptic-like activity simulated in 14 synchronous cortical areas of smaller extent (Scenario 3ter). **A. Top.** Brain maps of simulated activity (ground truth) and of raw activity computed with the three methods, normalized by their maximum. Positively polarized activity appears in shades of red, and negatively polarized activity in shades of blue. **Bottom.** Brain maps of activity after thresholding. **B.** Boxplots displaying sensitivity, precision, DLE and distance between distributions of activity by regions, with all methods and for all 20 instances. **C.** Heatmap showing average normalized activity by region for ground truth and the three methods.

### 2.2. Real HR-EEG data

Previous experimental results on simulated EEG data demonstrated that the APESISSY method performs consistently better than wMNE and sLORETA across all scenarios. To further verify whether APESISSY can maintain its good behavior on real data, we propose to qualitatively assess its efficiency on real ictal HR-EEG data from a patient with typical CAE (age 8, sex F, medication lamotrigine). An oriented lead field matrix of size 15002×256 was obtained following the same process we did for simulated data, using the BEM (Gramfort et al. 2010) with a 3-layer model comprising the inner skull, outer skull and scalp (conductivities 1, 0.033 and 1 reciprocally) after deriving a 15,002-vertex cortex mesh from the patient’s MRI-registrated brain. Bandpass filtering between 1 and 100 Hz, Notch filtering at 50 Hz, and elimination of bad channels followed by spatial interpolation were applied to the EEG data before source localization algorithms. In addition, we also applied spatial filtering using Independent Component Analysis (ICA) to remove electrophysiological artifacts (eye movements, electrocardiogram, …), as well as noise seemingly unrelated to the seizure. As shown in Fig. 8, we selected three time points – corresponding to the peaks of spikes occurring at the onset, middle and end of the seizure – at which the source imaging methods were applied.

**Figure 8.**
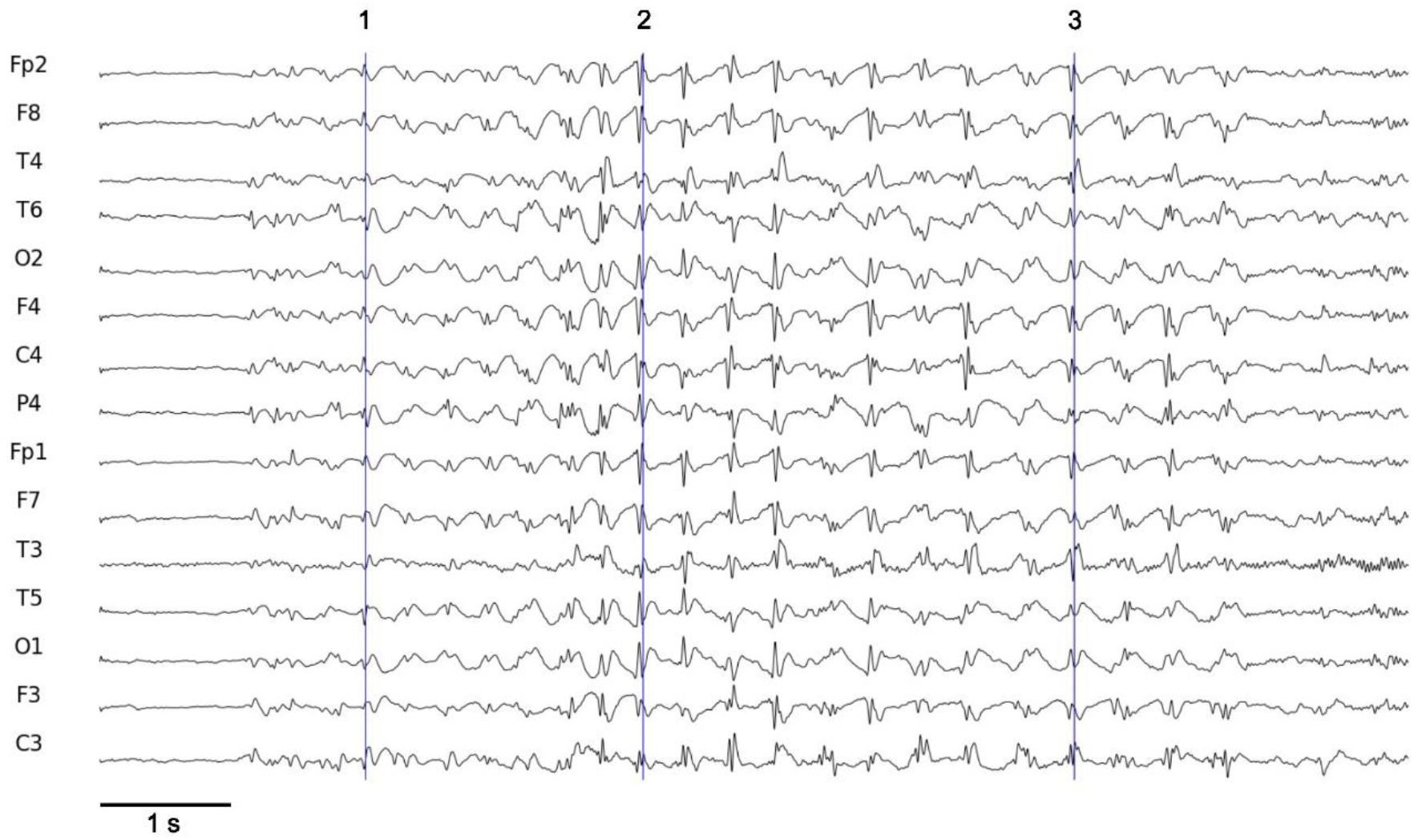
Real EEG signal displaying a typical spike-wave discharge on a few selected channels. Blue vertical lines mark the instants selected for source localization.

Fig. 9 A depicts, for each time point, the source localization results obtained with different algorithms after thresholding (40% of the maximum activity in absolute value). Fig. 9 B was obtained by summing, for each method and time point, the binarized activity of the dipoles composing the Desikan-Killiany atlas regions. The activity per region was then normalized between 0 and 1. It is interesting to see that the three methods reveal a significant temporal component of the seizure, with in particular a consensus on the involvement of the inferior temporal, fusiform and parahippocampal gyri, especially at the beginning of the seizure. However, temporal activity is not described identically by the three methods. Indeed, sLORETA detects some activity in the lingual gyrus, while APESISSY and wMNE detect the middle temporal gyrus. Both sLORETA and wMNE detect some activity in the entorhinal gyrus as well. The three methods also find extensive frontal activity associated with the second and third spikes, especially in prefrontal and orbitofrontal regions. However, prefrontal activity is limited to RMF for APESISSY – and detected as soon as the first spike –, while it extends to SF for sLORETA and wMNE. Apart from these two temporal and frontal main components, sLORETA detects some activity in the cingulate cortex, both wMNE and APESISSY detect some small active patches in the superior parietal cortex, and wMNE detects some activity in the right lateral occipital cortex.

**Figure 9.**
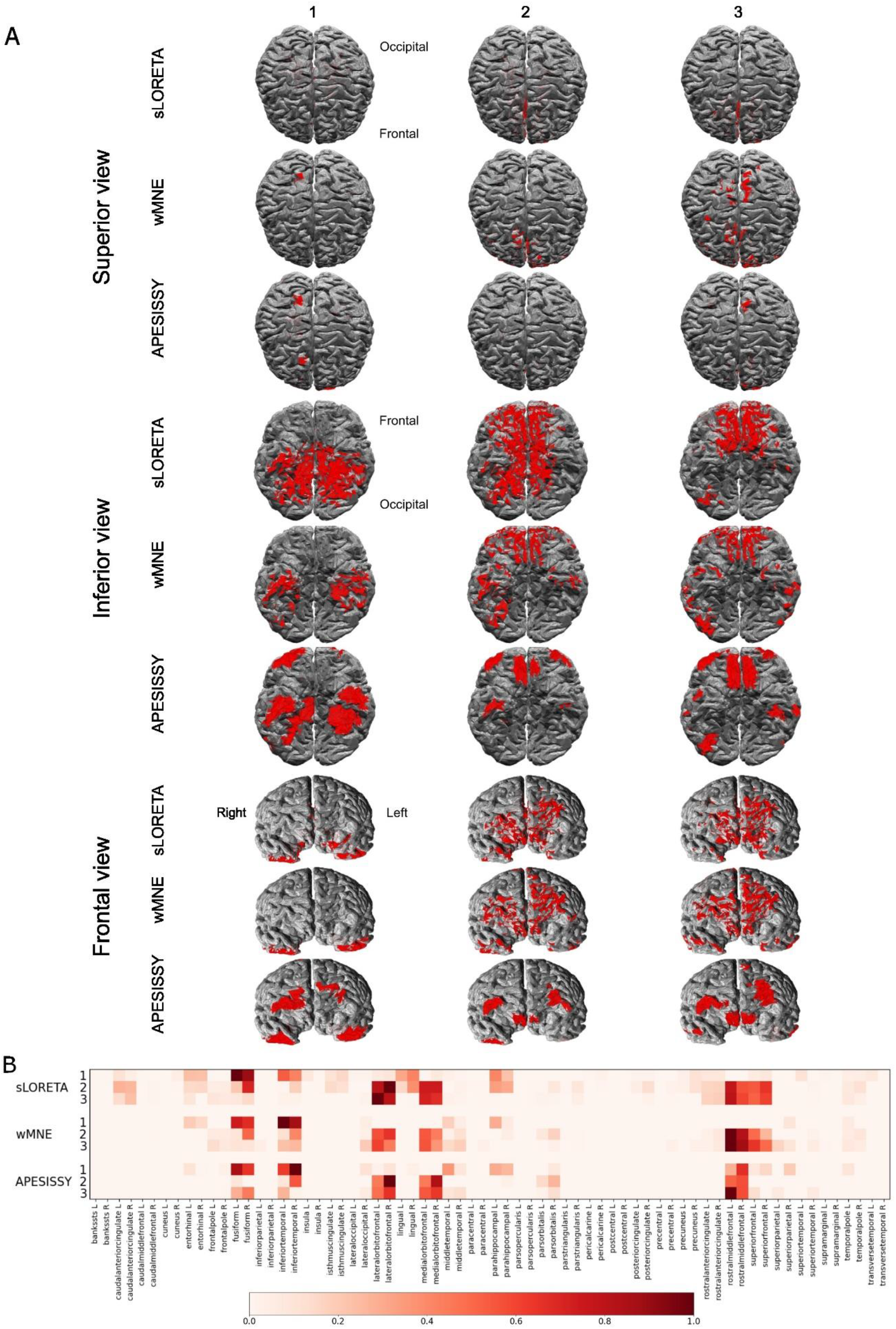
Comparison of methods for source localization on real high density EEG data capturing a typical absence seizure. **A.** Projections of brain maps of activity (in red) after thresholding for three selected instants (left to right) and according to the three methods. **B.** Time course heatmaps of normalized regional activity. For each method, the source localization at the three selected instants is displayed vertically from top to bottom. Activity is normalized relative to the highest percentage of activity at each instant.

Overall, as seen in simulated data, localized sources with wMNE and APESISSY are more circumscribed than sources found with sLORETA, especially those of APESISSY. Regions of activity rebuilt with APESISSY are also more cohesive and more clearly defined than those rebuilt by the two other methods.

## 3. Discussion

With APESISSY, we aimed to improve the localization of focal and multifocal distributed epileptic sources. Two parameters were considered when designing our simulations: the number of synchronous epileptic sources and the extent of these sources. It is worth noting that, in this context, the conventional ESI methods, such as wMNE and sLORETA, suffer from limitations, which make their solutions difficult to interpret in practice. Indeed, even if wMNE generally targets well the EZ, its estimated source localizations are often blurred. sLORETA also provides blurred solutions and tends to identify a large number of small brain regions. A further limitation with the regularized least squares-based ESI methods lies in determining the optimal regularization parameter value. To cope with all these limitations a new regularized least squares approach, named APESISSY, which can be seen as an improvement of our SISSY method, is proposed. In the same spirit as SISSY, this new algorithm considers the sparsity of both the sources and their spatial gradient. The big difference lies in the fact that the regularization parameter, which balances between the amount of data fidelity and regularization, is identified automatically. This makes APESISSY more convenient, especially in clinical routine. To assess the behavior of the proposed algorithm, a study on realistic simulated HR-EEG data, where ground-truth information is accessible, was conducted under different scenarios. An example on real HR-EEG data was also provided to validate the feasibility and the reliability of APESISSY on a clinical setting. A systematic comparison with wMNE and sLORETA was also provided.

### Simulated data

The performance of the three methods was assessed using four statistical indices after binarization of the dipoles’ activities through a thresholding step. Sensitivity measures how much the original activity is preserved in the reconstruction, while Precision measures how much artefactual activity is introduced by the used ESI method. Regarding the DLE, it measures the spatial accuracy of the solution compared to the original distribution of activity. Finally, the fourth index, the DD metric, is an atlas-based index assessing how close the solution is to the ground truth regarding its distribution of activity by regions.

The visual inspection of the results shows that APESISSY generally succeeds in localizing the targeted sources and the obtained solutions are more circumscribed for all conducted experiments, conversely to wMNE and sLORETA which to a large extent result in blurred solutions. This visual perception is confirmed through our quantitative evaluation. Indeed, APESISSY seems more efficient than wMNE and sLORETA, whatever the studied scenario and the assessed metrics. More precisely, APESISSY demonstrated the highest Sensitivity and Precision, with sLORETA and wMNE ranking second for Sensitivity and Precision, respectively. The Sensitivity gap between wMNE and APESISSY was at its most pronounced for scenarios involving large sources but remained significant with sources of lesser extent. The Precision gap between sLORETA and APESISSY was considerable for all scenarios, while this gap is minimized between wMNE and APESISSY in the case of epileptic sources with smaller extent. Furthermore, APESISSY also consistently exhibited significantly lower DLE than the other methods, particularly in scenarios involving a limited number of large-extent sources. When mapping activity to regions, APESISSY displayed superior accuracy as well, with the sole exception of scenario 3ter involving sources of small extent, for which wMNE exhibited slightly inferior index values, but the difference was not statistically significant.

Interestingly, the extent of sources had more impact on performance than the number of sources (Fig. 6-7). In general, the methods demonstrated superior performance in accurately locating and delineating activity patches when sources were of small extent (20/100 dipoles). However, although in these scenarios the three methods provided an adequate reconstruction of superficial sources, none of them could detect the “deep” sources PCUN and PSTC. This is in conformity with what has been reported concerning sLORETA (Wagner et al. 2004): when superficial and deep sources are simultaneously active, the latter tend to go unnoticed. In the case of large sources, regardless of the number of sources, wMNE and sLORETA mostly failed to capture the actual spatial distribution of activity, displaying inhomogeneous and blurred activity. Both methods detect occasional ghost sources sometimes located relatively far away from the targeted sources. In contrast, APESISSY is able to provide more robust solutions, especially in the cases of limited number of sources.

Consequently, APESISSY appears to be a robust alternative to commonly used methods such as sLORETA and wMNE. In particular, its advantage in terms of combined Sensitivity, Precision and DLE, demonstrates its superior ability to accurately assess both the location and extent of one “active” region. This makes it more reliable method if the purpose is to delineate the EZ in focal epilepsies. This is especially true given that we ignore a priori the extent of the EZ, which can be relatively large, and that APESISSY’s accuracy remains satisfactory in cases of extensive areas of activity, unlike the two other methods. Now, if the purpose is to localize multiple EZ, APESISSY can prove a valuable method as well. In fact, although it shares with wMNE and sLORETA some limitations, especially when it comes to the restitution of deep sources, APESISSY shows several advantages over its competitors. Apart from its better general ability to deal with extended sources, it equally proves more efficient in delineating and separating close sources. In addition, although its significant superiority over wMNE in terms of sensitivity does not always translate into a better capture of the spatial distribution of sources at the scale of the brain, it reflects a more accurate local mapping of sources’ activity.

An analysis of the raw values of current density provides some insight into the performance discrepancies between APESISSY and the two other methods. As the simulations only involved positively polarized current densities, any negativity found in the rebuilt sources is artefactual. The solutions provided by sLORETA and wMNE display two distinct types of negativities: local negativities within the boundaries of the actual sources, whose geometry usually coincides with that of the local sulci, and large-extent ones that occur in the vicinity of the targeted sources, potentially covering large parts of the cortex. In contrast, APESISSY’s solutions do not display negativities of the first type, and those of the second type are more circumscribed and of lower amplitudes. The first type of negativities involves gradients of amplitude of current source density within the boundaries of the actual sources. These gradients account for the inhomogeneities of the patches rebuilt with sLORETA and wMNE, and result in a loss in Sensitivity. The second type of negativities accounts for the majority of ghost sources, and results in a loss in Precision. We posit that two primary factors contribute to this differential handling of negativities. The first factor is the use of a spatial gradient regularization term in the APESISSY algorithm, which contributes to the spatial homogenization of the sources and the elimination of the first type of negativities. The second reason can be attributed to the L2 norm regularization term in sLORETA and wMNE. The L2 norm, which consists in the summation of the square values of the data points, tends to curb, more than the L1 norm, the maximal amplitude a dipole can attain. The contribution to the scalp EEG of a single dipole may then be compensated by that of multiple dipoles of lower amplitude. Indeed, in the cortex mesh, dipoles are characterized not only by their location, but also by their orientation, which is of great importance as the contribution of a dipole to the EEG is determined by the scalar product between its orientation and the vectors linking its location to the electrodes’ locations. When an active region contains sulci and gyri, the solution will change the polarity of some of the positively polarized dipoles in the ground truth, based on their orientation, in order to generate the same signal at the scalp level while keeping the dipoles’ amplitude lower, resulting in negativities of the first type. Similarly, the solution will potentially incorporate negatively polarized dipoles at some distance from the ground truth sources, resulting in negativities of the second type.

Note that, in our study, all simulations involved almost perfectly synchronized sources exhibiting same polarity at the chosen instant for source localization. This may constitute a limitation as real EEG data may typically derive from multiple active areas with mixed polarities, even in multifocal bilateral synchronous epilepsies seizures, for which synchrony between signals on different channels can be imperfect (Fig. 10).

**Figure 10:**
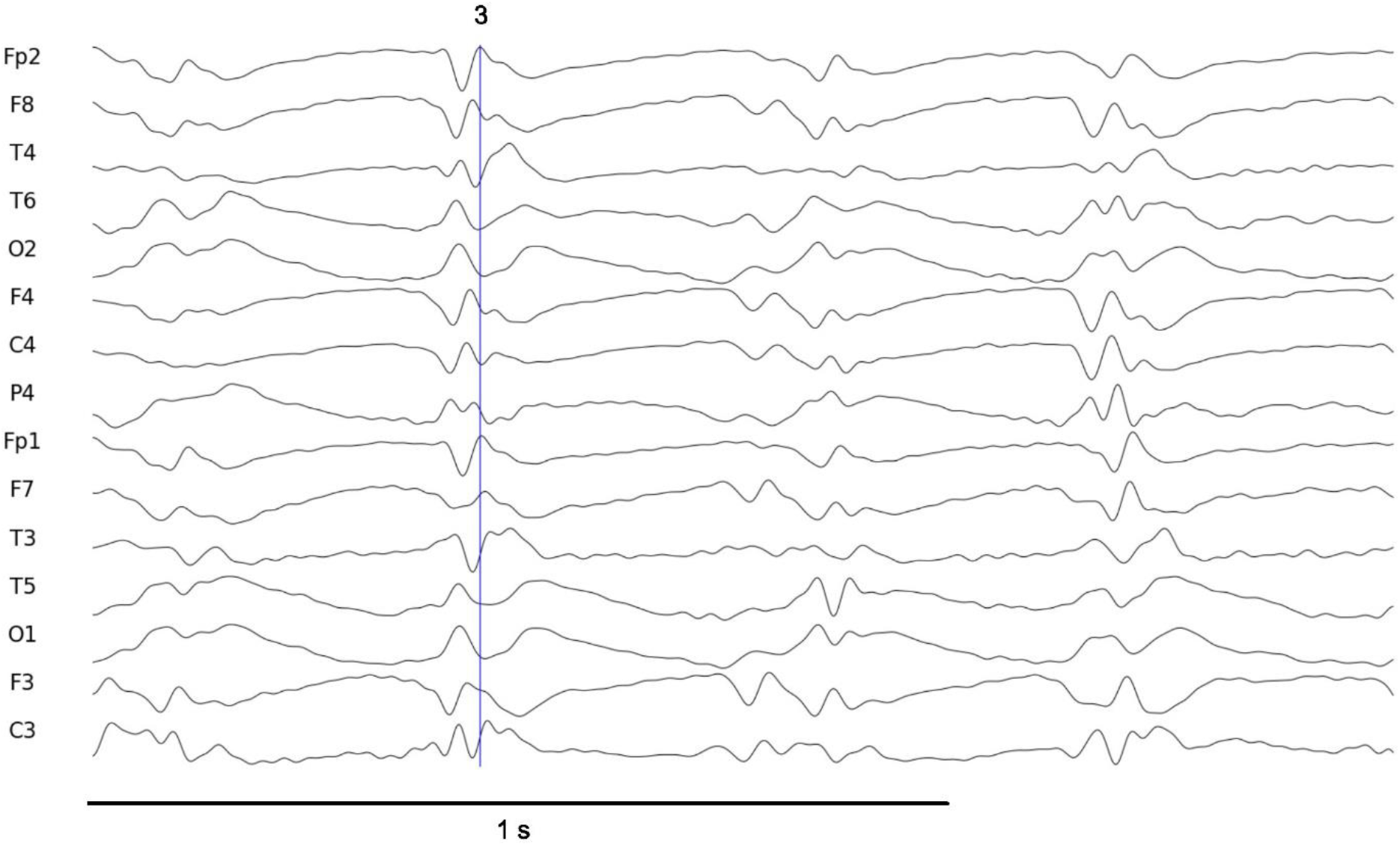
Real EEG signal displaying part of a typical spike-wave discharge on a few selected channels. Note the imperfect synchrony between the different channels. The blue vertical line marks the instant selected for source localization.

### Clinical HR-EEG data

When applied to real HR-EEG data from a patient experiencing a typical CAE seizure, the three methods agreed on the involvement of temporal and frontal regions. The prefrontal and orbitofrontal involvement is well documented through fMRI studies (Moeller et al. 2008; Li et al. 2009; Bai et al. 2010; Berman et al. 2010; Masterton et al. 2013; Carney and Jackson 2014), MEG (Westmijse et al. 2009; Tenney et al. 2013; Miao et al. 2014; Wu et al. 2017; Gadad et al. 2018; Wang et al. 2023) and EEG (Holmes et al. 2004; Tucker et al. 2007; Sarrigiannis et al. 2018; Jun et al. 2019). However, in the studied case, no used ESI method detected extensive parietal activity, although both the parietal cortex (Moeller et al. 2008; Bai et al. 2010; Berman et al. 2010; Tenney et al. 2013) or the temporo-parieto-occipital region are frequently associated with CAE seizures (Tang et al. 2016; Wu et al. 2017). Interestingly, the detection of temporal and more generally ventral activities is consistent with findings from other EEG (Yeom et al. 2015; Sarrigiannis et al. 2018) or fMRI studies (Bai et al. 2010). However, it should be noted that part of this activity, particularly that associated with the first spike, might be artefactual. Indeed, when localizing sources from real EEG data, two new potential sources of error arise compared to simulated data: non-physiological noise and artifacts, on one hand, and bias associated with the head model or lead field matrix, on the other hand. Although artifacts were mitigated using ICA, we cannot dismiss the possibility of a bias introduced by the lead field matrix. A slight overestimation of the signal attenuation from some regions could result in an inflated estimation of their activity, as computed by the ESI methods. Finally, as expected from simulated data, none of the three methods detected any deep source, in contrast to fMRI and MEG methods regularly reporting PCUN and PSTC activity during epileptic absences (Moeller et al. 2008; Li et al. 2009; Masterton et al. 2013; Miao et al. 2014).

When comparing the ESI methods, three main points can be highlighted. First, as shown in simulated experiments, APESISSY provides more homogeneous and better delineated source localizations, even for fairly close regions, compared wMNE and sLORETA. This suggests that APESISSY may yield results that are more interpretable and clinically relevant in practical situations. Secondly,sLORETA seems to detect more widespread ventral activity, especially in temporal areas, than wMNE and APESISSY. It also detected some activity in the cingulate cortex. However, since sLORETA was prone to artefactual activation in the cingulate cortex during simulations, we suspect this finding could be artefactual as well. Thirdly, aside from the SF region and the right lateral occipital cortex, wMNE and APESISSY generally converge in detecting similar active regions. This convergence aligns with the observation between these two methods with simulated data involving patches of activity of a small extent.

As mentioned earlier, localizing sources from real EEG data constitutes a distinct challenge compared to using simulated data. In particular, as illustrated in Fig. 10, there is no strict synchrony between spike-waves across different channels. Therefore, selecting a specific time point involves a degree of arbitrariness, which can introduce biases that may significantly impact the sources retrieved. Since the aim here was to compare ESI methods, we employed the same point-in-time approach used with simulated data. However, to achieve a more accurate representation of the regions involved, we recommend adopting statistical approaches based on time intervals.

## 4. Conclusion

This paper introduces APESISSY, a novel regularized least squares ESI method imposing double sparsity on the spatial gradient of current density and the current density itself. Contrary to the original SISSY algorithm, the regularization parameter is identified automatically, making APESISSY more practical for clinical use, and paving the way for close-loop EZ localization. The use of realistic simulated HR-EEG data showcased that, when compared to state-of-the-art methods, APESISSY is very efficient in accurately localizing EZ and reconstituting homogeneous well-delineated areas of activity. Notably, APESISSY is effective even in the presence of extended sources, which makes it particularly valuable in determining the spatial distributions of sources in cases of both unifocal and multifocal epilepsies. In addition, compared to wMNE and sLORETA, APESISSY seems to be the most promising approach to deal with synchronous epileptic patch configurations, an especially challenging scenario for source imaging. The behavior of APESISSY was also qualitatively assessed on real HR-resolution EEG data obtained from a patient with typical CAE. Given its interesting capabilities, APESISSY looks promising in clinical EEG. All these aspects should be further investigated in the future by evaluating the proposed approach on broader clinical data in order to confirm the preliminary results of this paper.

## Acknowledgements

This study has been funded by the Institut des Neurosciences Cliniques de Rennes (INCR, https://www.incr.fr), as part of the PREDILEPSY Project.

## Annex

APESISSY pseudocode

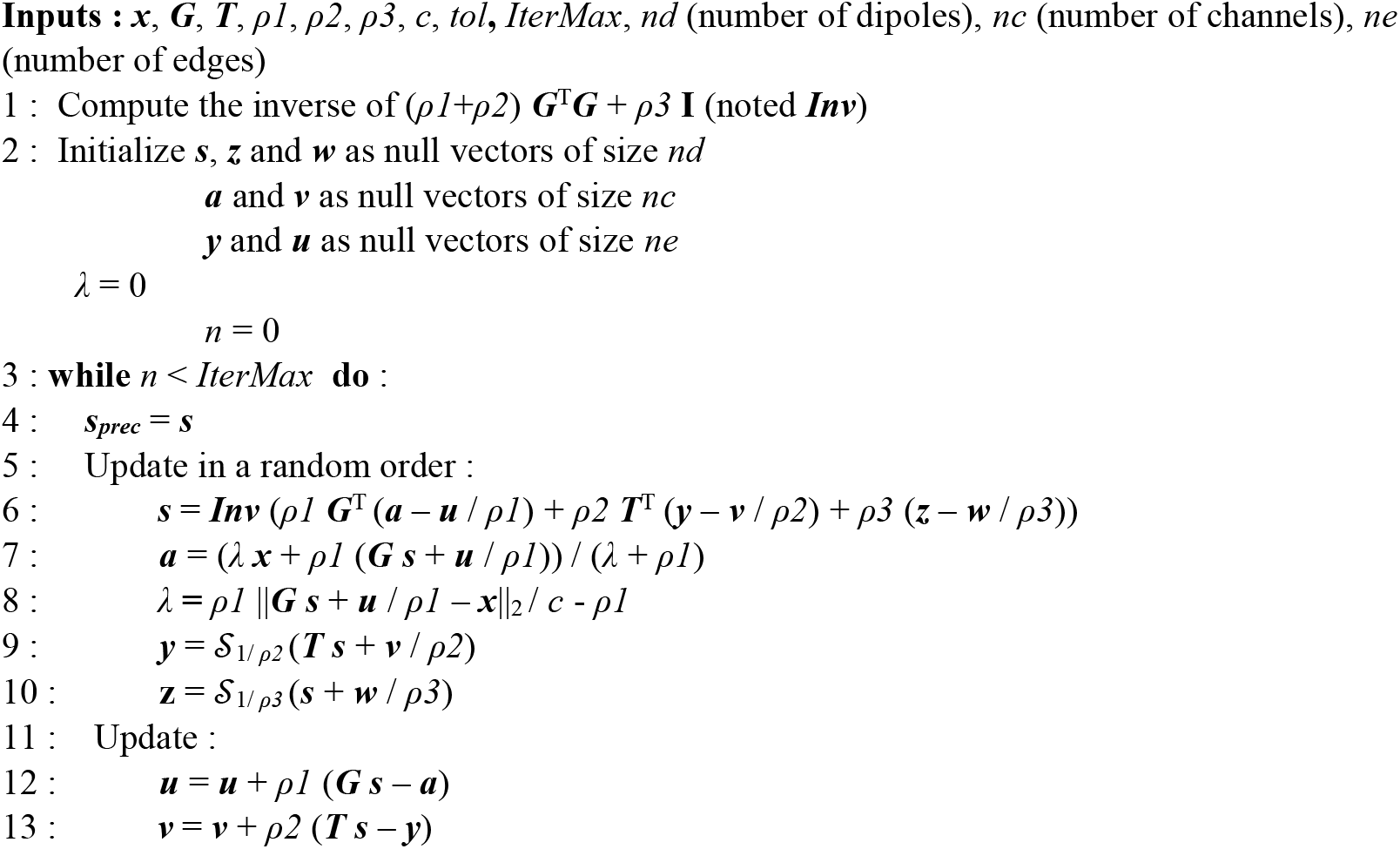
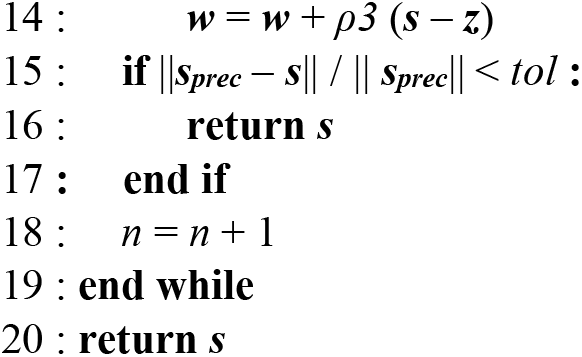

